# Synergistic Antimicrobial Activity of Epinecidin-1 and Its Variants with Antibiotics Against Nosocomial Pathogens: Implications for In Vivo Wound Healing

**DOI:** 10.1101/2025.07.02.662646

**Authors:** Sivakumar Jeyarajan, Anbarasu Kumarasamy

**Affiliations:** Microbial Biotechnology Laboratory, Department of Marine Biotechnology, Bharathidasan University, Tiruchirappalli, Tamil Nadu, India; Transgenic Animal Model Core, Biomedical Research Core Facilities, University of Michigan, Ann Arbor, Michigan, USA

**Keywords:** Antimicrobial peptides, Epinecidin-1, Multi drug resistance, Nosocomial, Wound healing, Antibacterial

## Abstract

In this study, we investigated the antimicrobial properties of the antimicrobial peptide (AMP) epinecidin-1 and its variants against a range of nosocomial bacterial pathogens. The bacteriostatic effects on *Escherichia coli, Haemophilus influenzae, Klebsiella pneumoniae, Pseudomonas aeruginosa, Streptococcus pneumoniae*, and *Staphylococcus aureus* were evaluated using the microbroth dilution technique. Mechanistic insights into the antimicrobial action were studied through acridine orange and ethidium bromide (AO/EtBr) staining, which revealed the ability of AMPs to form pores in bacterial membranes. Furthermore, the wound healing efficacy of these peptides was assessed in vivo using rat model, where they significantly accelerated the healing process. Following the identification of their bacteriostatic activity against pathogenic strains, we also examined the effects of combining these AMPs with conventional antibiotics, namely ampicillin, kanamycin, and vancomycin. Epinecidin-1 and its variants displayed synergistic and additive interactions with these antibiotics at certain concentrations. This combination approach is particularly promising for treating infections in both human and livestock populations, thereby potentially mitigating the risk of environmental pathogen outbreaks and complications of multidrug resistance. In summary, our findings suggest that epinecidin-1 and its variants not only exhibit potent antimicrobial and wound healing properties but also enhance the efficacy of traditional antibiotics. These characteristics make them strong candidates for clinical and industrial applications aimed at managing bacterial pathogenicity.

## 1. Introduction

Antibiotics had been promising therapy against pathogenic infections, however, their utility is rapidly declining due to the growing threat of antibiotic resistance—a problem intensified by poor management practices and a lack of progress in developing new antibiotic classes. With the global exploitation of antibiotics, the quick spread of bacteria resistant to antibiotics has been growing year after year [1]. Especially nosocomial bacterial resistance to antibiotics originating from hospitals are a major threat. The nosocomial pathogens like *Enterobacter* species, *Staphylococcus aureus, Klebsiella pneumoniae, Pseudomonas aeruginosa*, cause the majority of hospital infections with higher economic burden in the health care sector, morbidity and mortality to the patients.

These pathogens are able to evade the effects of commonly prescribed antibiotics and are often linked to a substantial economic impact, primarily by prolonging hospital stays and reducing overall workforce productivity. To overcome antibiotic resistance, 10 to 100 times more quantities of antibiotics were prescribed. However this did not offer any cure rather it led to resistance and deteriorated the intestine, metabolism and excretory organs of humans. This makes antibiotic resistance one of the greatest threats to global public health in the 21st century as antibiotics steadily lose efficacy in clinical and community settings [2]. The World Health Organization has estimated that if the current pace of antibiotic resistance in medical treatment continues unchecked, it could result in 10 million fatalities yearly and a 2–3.5% annual decrease in gross domestic product by 2050 [3].

Addressing the escalating threat of antibiotic resistance necessitates not only the development of novel antimicrobial agents but also the implementation of innovative therapeutic strategies. These include the repurposing of preclinical compounds and the broader application of combination therapies to enhance efficacy and circumvent resistance mechanisms. Mechanistically, bacterial pathogens employ several strategies to evade antibiotic action. These include active expelling of antibiotics *via* membrane-bound efflux pumps [4], reduced permeability through the downregulation or modification of porins [5] and selective transporters [6], enzymatic degradation of antibiotics, and target modification through mutation or synthesis of alternative binding proteins [7]. To counteract these resistance mechanisms, preserving the intracellular concentration, identifying alternative membrane penetrating methods for quick entry and structural integrity of antibiotics or drugs are critical. Preventing efflux and enzymatic degradation can prolong the intracellular residence time of the drug, thereby enhancing its pharmacodynamic effect. In this context, small molecules and short-chain antimicrobial peptides have shown promise in stabilizing antibiotic activity and facilitating intracellular retention by inhibiting efflux pumps [8].

Antimicrobial peptides (AMPs), whether naturally occurring or synthetically engineered, represent promising alternatives to conventional antibiotics [9] [10]. These peptides are typically rich in hydrophobic amino acids and possess a net positive charge, enabling them to interact with and disrupt negatively charged microbial membranes.

Their primary mode of action involves neutralizing bacterial surface charges, permeabilizing membranes, and ultimately inducing cell lysis, which significantly reduces the likelihood of resistance development. Compared to traditional antibiotics, AMPs are generally regarded as safer, with a lower propensity for adverse side effects and resistance emergence. They exhibit broad-spectrum activity against a wide range of pathogens, including bacteria, fungi, protozoa, and also demonstrate antiviral and anticancer properties. In addition to their intrinsic antimicrobial activity, small molecules and short AMPs can enhance antibiotic efficacy through several mechanisms: (a) acting as carriers to facilitate antibiotic entry into microbial cells [11]and prevent efflux, (b) acting as adjuvants to prevent antibiotic degradation [12], and (c) inhibiting degradative enzymes by occupying their active sites [13-15], thereby preserving antibiotic stability.

Given the multifaceted roles of short antimicrobial peptides (AMPs), this study evaluated their potential synergistic effects when used in combination with conventional antibiotics. To assess this, we tested AMPs alongside three distinct classes of antibiotics: ampicillin, kanamycin, and vancomycin. Kanamycin, an aminoglycoside antibiotic, exerts its bactericidal effect by binding to the 30S ribosomal subunit, thereby disrupting protein synthesis. It is particularly effective against Gram-negative bacteria and is commonly employed in both clinical settings and laboratory research, especially for treating infections resistant to other antibiotics. Ampicillin, a β-lactam antibiotic within the aminopenicillin subclass, inhibits bacterial cell wall synthesis by targeting penicillin-binding proteins (PBPs)[16]. It exhibits broad-spectrum activity against numerous Gram-positive and selective Gram-negative organisms and is frequently used to treat respiratory, urinary tract, and gastrointestinal infections. Vancomycin, a glycopeptide antibiotic, also targets cell wall synthesis but does so by binding to the D-Ala-D-Ala terminus of peptidoglycan precursors [17]. This mechanism makes it highly effective against

Gram-positive bacteria, including resistant strains such as MRSA. Due to its potential adverse effects, including nephrotoxicity and infusion-related reactions like “red man syndrome,” vancomycin is typically reserved for severe infections.

In previous work, we characterized the antibacterial activity of epinecidin-1 and its variants [18-20]. In the current study, we investigated whether combining these peptides with non-peptide antibiotics could enhance antimicrobial efficacy and achieve complete inhibition of bacterial growth. Beyond assessing antimicrobial potency, we also explored the underlying mechanisms of action and evaluated the wound-healing properties of the peptides.

## 2. Materials and Methods

### 2.1. Peptides

Antimicrobial peptides (AMPs) epinecidin-1 (epi-1) and its variants namely var-1 and var-2 (purity ≥ 95%) were using 9-fluorenylmethoxycarbonyl (Fmoc) solid-phase peptide synthesizer by GenicBio Limited, Shanghai, China. AMPs sequences as reported in (ref) with their corresponding lysine replaced sites are shown in Table 1.

**Table 1.** Sequence of epinecidin-1 and its variants. Lysine replaced residues are highlighted, bold, italicized, and underlined.

| Peptide name | Sequence |
| --- | --- |
| Epinecidin-1 | GFIFHIIKGLFHAGKMIHGLV |
| Variant 1- Replacement of H with K | *GFIF<u>K</u>IIKGLF<u>K</u>AGKMI<u>K</u>GLV |
| Variant 2- Replacement of A with K | *GFIFKIIKGLF<u>K</u>GKMIKGLV |

\*- Replaced residues are represented in bold letters and underlined.

### 2.2. Peptide Characterization by Reverse Phase-High Performance Liquid Chromatography and Structural Assessment

The peptides were passed through Eclipse Plus C18 (Agilent Technologies, Santa Clara, CA, USA) reverse phase column (4.6 mm × 150 mm, pore size 5 µm) and eluted by a linear gradient of 0%–100% (v/v) acetonitrile in water containing 0.1% (v/v) Trifluoroacetic acid at a flow rate of 1.0 mL min−1, by monitoring the absorption at 280 and 214 nm UV detectors.

For structural assessment, hydrophobic moment, electrostatic surface and Ramachandran plot; secondary structure parameters were performed. Hydrophobic moment: a measure of amphiphilicity which aids to partition in the membrane was calculated using Eisenberg consensus scale available in HELIQUEST server [21]. For plotting electrostatic surface map and Ramachandran map, structural coordinates were required.

Hence the structural coordinates were obtained from i-TASSER (Iterative Implementation of the Threading ASSEmbly Refinement) [22]. From the structural coordinates, electrostatic surface maps were generated using PyMOL (Molecular Graphics System, Version 2.5.2 Schrödinger, LLC (New York, NY, USA). The Ramachandran map secondary structure details (phi/psi) plots for the peptides were calculated using RAMPAGE tool [23, 24] and VADAR[25].

### 2.3. Assessment of Antibacterial Activity of Epinecidin-1 and its variants

Gram negative; *Escherichia coli* (*E. coli* MTCC 2939), *Pseudomonas aeruginosa* (*P. aeruginosa* MTCC 741), *Haemophilus influenza* (*H. influenzae* ATCC 25922), *Klebsiella pneumonia* (*K. pnuemoniae* MTCC 432) and Gram positive; *Staphylococcus aureus* (*S. aureus* ATCC 25923) and *Streptococcus pneumonia* (*S. pneumoniae* ATCC 49619) were selected as the test pathogenic microorganisms. The selected microorganisms were cultured in Mueller Hinton broth. Minimum inhibitory concentration (MIC) against bacteria were determined using 96-well microtiter cell culture plates in Mueller Hington broth as described earlier [26]. Briefly, bacterial cells grown overnight were diluted in medium broth to a cell density of 10^6^ colony-forming units (CFU) ml^-1^. This suspension was aliquoted to 100 µl in each well in a microtiter plate along with plain broth for sterility control and only suspension without peptide for growth control. Other than controls, 100 µl of peptide was dissolved in phosphate-buffered saline (PBS) corresponding to a concentration of 1024 µg ml^-1^, was mixed with 100 µl of suspension so that it equals to 512 µg ml^-1^. This peptide suspension was serially diluted to reach concentrations of 0, 2, 4, 8, 16, 32, 64, 128, 256 and 512 µg ml^-1^. The plates were incubated at 37°C for 16 h in an incubator.

After incubation, bacterial growth was measured by absorbance readings (optical density) at 595 nm using a microtiter plate reader (Bio-Rad, Hercules, CA, USA). From the O.D readings, the MIC value was determined. The MIC was defined as the lowest concentration of a peptide that inhibited growth of the bacteria after 16 h incubation. Each experiment was performed in triplicate with mean ± standard deviation (S.D). Tukey’s multiple comparisons test (two-way ANOVA) was performed for statistical analysis between epi-1 vs var-1 and epi-1 vs var-2. The statistical significance representations are a; p < 0.5, b; p < 0.1, c; p < 0.01, and d; p < 0.001.

### 2.4. Acridine Orange (AO) Ethidium Bromide (EtBr) staining

Gram negative; *Escherichia coli* (MTCC 2939), *Pseudomonas aeruginosa* (MTCC 741), *Klebsiella pneumonia* (MTCC 432) and Gram positive; *Staphylococcus aureus* (ATCC 25923) were selected as the test microorganisms to differentially stain live and dead cells with Acridine orange and Ethidium Bromide. Bacterial cells were grown overnight, diluted in the broth medium to a cell density of 10^6^ colony-forming units (CFU) ml^-1^. Bacterial strains were treated with the peptides at their respective MIC for 16 hours. Untreated cells were used as control. For AO/EtBr staining, the untreated and treated cells were centrifuged at 4000 x g for 10 minutes. The pellet was washed with PBS and stained with 100 µl of PBS containing 1 µg ml^-1^ AO and 1 µg ml^-1^ EtBr for 30 minutes. Excess dye was washed with PBS twice. The stained cells were subsequently observed, and pictures were taken under a fluorescence microscope (Leica inverted microscope, Deerfield, IL, USA) at 40X magnification.

### 2.5. Combined antibacterial effect of peptides with antibiotics

After finding the bacteriostatic effect of peptides, ability of peptides to act in combination with clinical antibiotics such as ampicillin, kanamycin and vancomycin was evaluated for antagonistic, additive or synergistic effect. Concentrations: 3.75, 7.5, 15 and 30 µg were used. Example 3.75 denotes 3.75 µg of peptide and 3.75 µg of antibiotic were used individually and in combination. Loewe additivity [27]was used to study the effect of peptide and antibiotics in combination. The combination index is calculated by the sum of ratios of concentrations of drugs in combination divided by concentration of drugs used individually. The formula for Combination index (CI) =(FAp+FAa)/

FAcombination. FAp represents Fraction affected by peptide; FAa represents fraction affected by antibiotic; and FAcombination represents fraction affected by combination of AMP and antibiotic at their individual concentration. The graphs showing the growth percentage of growth is shown in the supplementary. The effect of combination showing the combination index is shown in the main article. Combination index CI value <1 is Synergy, =1 is additive and >1 is antagonistic.

### 2.6. Wound healing assay

Male Wistar albino rats (∼150g) were used for this study according to Institutional Animal Ethical Committee guidelines (Ref. No. BDU/IAEC/2012/42) Bharathidasan University, Tiruchirapalli, India. The dorsal hairs were removed, and the skin was cleansed with PBS. Four equal thickness cutaneous wounds under aseptic conditions were created in the skin on the dorsal side of each rat; two on the left and two on the right, 1 cm away along the midline vertebral column. Skin biopsy punch of 5-mm-diameter was used. After wounding, rats were caged individually until termination of the experiment. Wounds were cleaned with PBS. Epi-1 and its variants at 50 µg/ml (dissolved in PBS) were applied directly to the wounded site twice daily. After application of peptide, petroleum jelly was applied on the wounded surface to sustain the peptide within the infected site for preventing the spread of peptides to other areas. Petroleum jelly alone applied to wounded animals served as vehicle control. Wound healing was macroscopically monitored by taking digital photographs after every two days. After 8 days, the animals were euthanized, and wound tissues were collected for histological studies. Wound tissue specimens were fixed in 10% neutral-buffered formalin solution and embedded with paraffin. The paraffin embedded wounded tissue was sliced to a thickness of 5 µm and stained with hematoxylin and eosin as described by [28].

## 3. Results

### 3.1. Characterization of epi-1 and its variants

Numerous studies, following extensive scrutiny and biological characterization, have provided evidence that enhancing the cationicity and α-helicity of antimicrobial peptides (AMPs) results in improved antimicrobial activity [18, 26, 29]. This enhancement can be achieved by incorporating lysine, which promotes helix formation, thereby increasing AMP activity [30]. To augment the cationicity and helicity, lysine was substituted into the template sequence of AMP epi-1 to create var-1 and var-2, as detailed in the materials and methods section. The biophysical properties of these peptides are presented in Table 2. With lysine substitution, the overall charge and isoelectric point (pI) of the peptides increase. The hydrophobic moment, a measure of amphipathicity, is higher for var-1 compared to var-2, as var-1 possesses a more perfect amphipathic structure. In contrast, var-2 contains an additional lysine residue in the hydrophobic region, leading to an increase in charge rather than amphipathicity. This property is also reflected in their reverse-phase high-performance liquid chromatography (RP-HPLC) profiles. The hydrophobicity of the peptides was assessed using RP-HPLC (Fig 1), which showed a strong correlation with their hydrophobic and hydrophilic characteristics based on retention times. The peptides were loaded on a RP-HPLC C18 column individually and eluted with a gradient of acetonitrile (0-100%) over 30 minutes (Supplementary Figure.1). The retention times are listed in Table 2. The normalized retention times relative to absorbance units for the three peptides are shown in Figure 1, with the elution order being var-2 (21.45 min), followed by epi-1 (23.7 min), and then var-1 (25.8 min). Based on the isoelectric point (pI), the expected elution order should be var-2 < var-1 < epi-1. However, lysine substitution affects the structural parameters. As a result, the elution profile for var-1 reflects its structural characteristics (hydrophobic moment and amphipathic nature) rather than its pI. Variant 1 was found to possess a more substantial hydrophobic patch (shown in next paragraph), which likely caused it to elute later than epi-1. Var-2, due to its greater hydrophobicity compared to Epinecidin-1, Var-2 eluted earlier at 21.45 minutes, whereas Epinecidin-1 eluted at 23.7 minutes during chromatographic separation.

**Table 2.** Biophysical characteristics of the AMPs.

| Peptide | Sequence | Molecular weight (Da) | Net Charge | pI | Hydrophobic Moment | Elution time (min) |
| --- | --- | --- | --- | --- | --- | --- |
| Epi-1 | GFIFHIIKGLFHAGKMIHGLV | 2335.8 | +2 | 10 | 0.432 | 23.7 |
| Var-1 | GFIFKIIKGLFKAGKMIKGLV | 2308.9 | +5 | 10.6 | 0.513 | 25.8 |
| Var-2 | GFIFKIIKGLFKKGKMIKGLV | 2366.0 | +6 | 10.7 | 0.477 | 21.45 |

**Figure 1.**
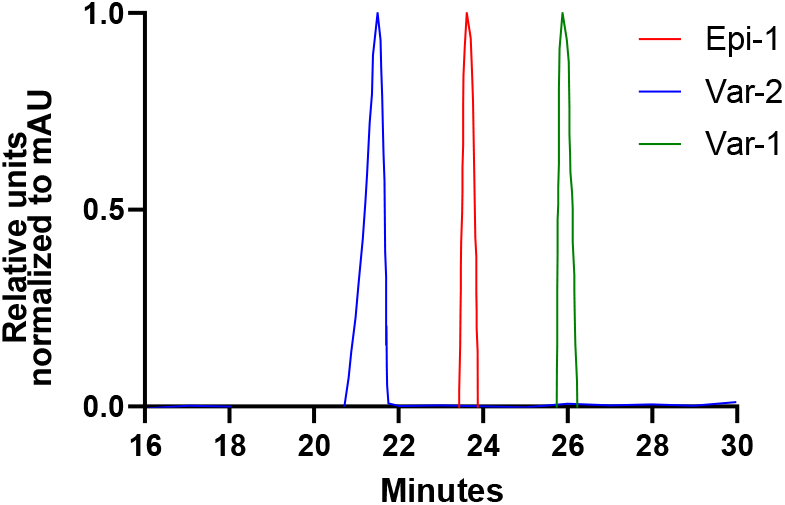
RP-HPLC profile of epi-1 and its variants. The peptides were loaded onto the C18 column and eluted with gradient increase of acetonitrile from 0-100% for 0-30 min. The increasing gradient scale of acetonitrile is given in the right Y-axis. The elution profile from 15 to 30 min of the normalized absorbance unit is shown in the figure and the individual elution profile is shown in the Supplementary Figure.1. Red, green and blue represent epi-1, var-1 and var-2. The elution order is var-2 (21.45 min), followed by epi-1 (23.7 min), and then var-1 (25.8 min). There is not a direct correlation of elution with the pI (iso electric point) however the elution profiles co-relate to the hydrophobic moment. This is because, var-1 has a perfect amphipathic structure, both hydrophilic and hydrophobic surfaces are equally distributed.

### 3.2. Electrostatic surface potential of epi-1 and its variants

The amino acid arrangement of the peptides when represented as a space-filled model with their electrostatic surface potential, the AMPs show an overall positively charged surface (red surface). In epi-1 (Fig 2a), the blue regions represent histidine, and the red region represents lysine. After histidine is substituted with lysine for var-1 (Fig 2b) the space-filled model transforms to a large hydrophobic (yellow) and positive charged hydrophilic (red) patch. A perfect amphipathic surface is seen. For var-2 (Fig 2c), additional substitution of lysine in the place of lysine, introduces a small hydrophilic area within the main hydrophobic region. This does not have a perfect amphipathic structure, but has large hydrophilic patch (red).

**Figure 2.**
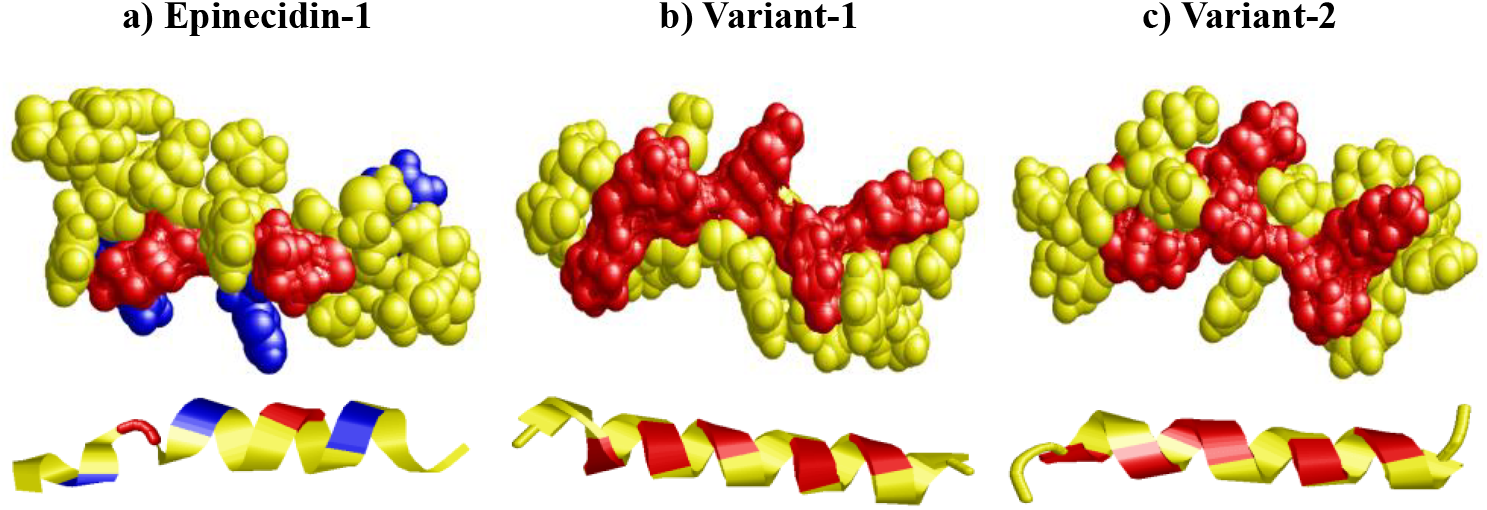
Electrostatic surface (top) and secondary structure (bottom) representation of epi-1 (a), var-1 (b) and var-2 (c). Histidine present in epi-1 is shown blue. For epi-1, the inherent lysine is shown red. Yellow represents amino acids other than histidine and lysine. For var-1 and var-2, both inherent and substituted lysine is shown red.

Below the space filled model, the ribbon diagrams of the peptides are shown. For epi-1, the first lysine does not form a helix. It rather forms a random coil. The histidine is shown as blue patches. Substitution of lysine at the areas of blue patch, transformed the coiled structure of epi-1 into helix structure in the variants. The variants do not have coiled structure.

### 3.2. Ramachandran plot of epi-1 and its variants

To validate and estimate the secondary structural characteristics, the phi and psi angles of the peptides were plotted in Ramachandran map quadrant space using RAMPAGE. Ramachandran map defining the distribution of amino acids in their corresponding geometric conformation featuring the favorable and allowed dimensions of the peptides by their steric-; intra-and inter-molecular forces are shown in Fig 3 a-c. This geometric structure in-turn reflects the stability of the peptides.

**Figure 3.**
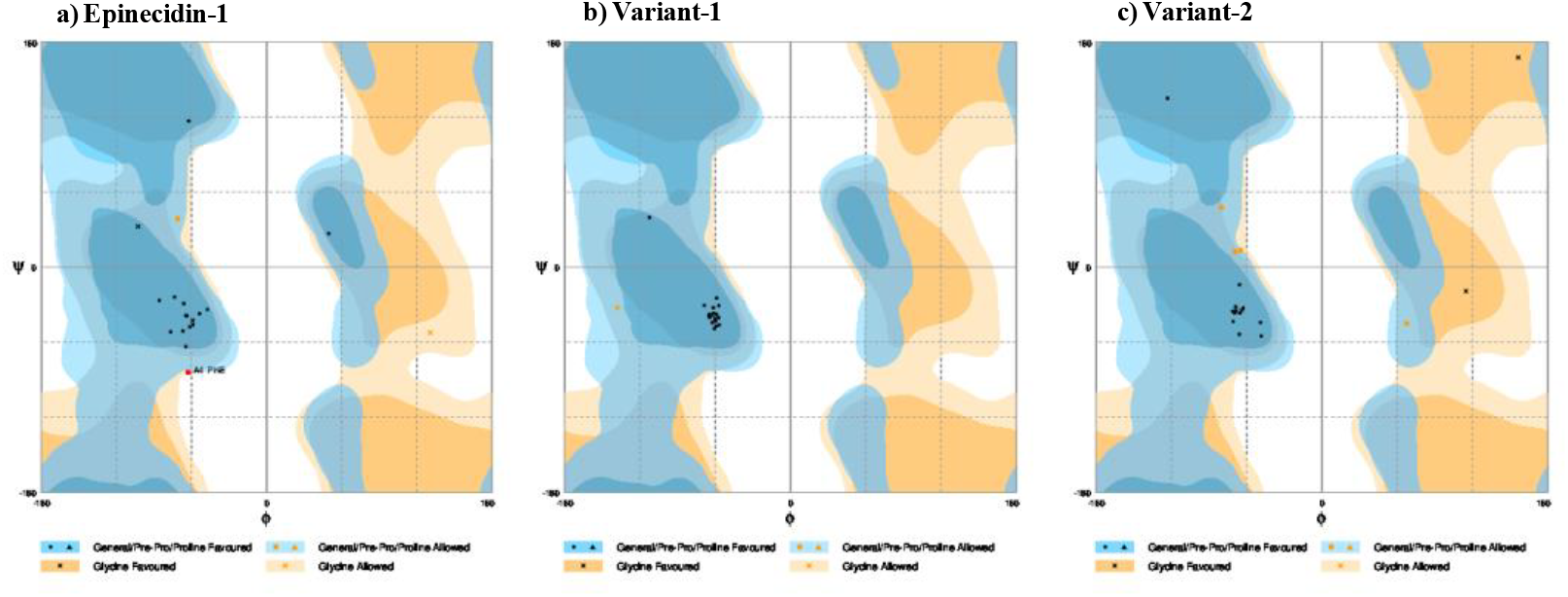
Ramachandran plot. The distribution of phi/psi angles of peptide bonds are shown for a) epi-1, b) var-1 and c) var-2. The figure depicts the thermodynamics, inter and intramolecular forces that stabilize the folded peptide structure. The Ramachandran plot plotted for each peptide is conglomerate of four types of geometrical arrangements (1) general: refers to 18 standard non-glycine non-proline amino acids, (2) glycine: plotted for glycine, (3) proline: plotted for proline and (4) pre-proline: residues that come before a proline. The plot is divided into regions; dark color represents “favored”, light color represents “allowed”, and white color represents “disallowed”. The spots represent the amino acids that fall on these regions based on steric conformation. The position of spots reveals the likelihood of finding a residue in that conformation. The general favored region and Pre-Pro favored region are indicated with dark blue color. The general allowed region and Pre-Pro allowed region is shown in pale blue. The Glycine favored and allowed regions are shown in dark and pale orange, respectively. The black squares (◼) represents general amino acids in the favored region. The orange squares 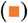 represents the general amino acids in the allowed region. The red squares 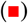 represent the general amino acids in the dis-allowed region. The glycine aminoacid is represented as “x”. Presence of “x” in the dark orange region represents glycine in the favored region and “x” in the pale orange represents glycine in the allowed region. The triangle represents the proline and pre-proline. However, for the peptides in this study there is no proline.

For epi-1 Fig 3a, the amino acids are distributed in the right-handed alpha helix region (psi angle between-60° to 0° and phi angle between-120° to 0°). One general amino acid fall in the allowed region and the phenylalanine in the fourth position falls in the disallowed region. For the four glycine aminoacids of epi-1, one glycine falls in the allowed region while the three other glycine falls in the favored region. For var-1, all the aminoacids are more confined in the core region of the perfect helix. Lysine substitution has mediated all the four glycine of var-1 to fall within the helical region. The psi angles for the var-1 fall perfectly between –60 to-30 and the phi angle falls between-30° to-80° which defines that the structure is confined to core of helix with energetically stable conformation (Fig 3b). For var-2, the aminoacids are distributed in the right-handed alpha helix region like epi-1. The glycine falls in the favored region. However, four aminoacids falls in the general allowed region. This shows var-1 is a very stable perfect helix compared to epi-1 and var-2. For the designed variants, all the glycine falls in the allowed region. This indicates that adding lysine guide the peptide to fold into a stable conformation. The steric conformation from the Ramachandran plot perfectly aligns with the amphipathicity predictions and the RP-HPLC profile.

The boundary marked by red represents the beta sheet and marked by yellow represents the right-handed alpha helix. All the peptide conformation falls in the alpha helix region. For epi-1 and var-2, couple of amino acids fall in the beta sheet and glycine favored regions. For var-1, all the amino acids fall in the perfect alpha helical core region with the maximum score.

After determining the steric conformation from the Ramachandran map, the quantitative secondary structural characteristics (percentages of helicity, beta sheet, and random coil) of the peptides were calculated using VADAR. Table 3 shows the percentages of these secondary structural elements and their distribution across favored, allowed, and outlier regions. The helical content of epi-1 is 47%, while var-1 and var-2 have 76%. The variants have a random coil content of 23%, reduced from the 52% seen in epi-1.

**Table 3.** Secondary structure quantitation statistics of the peptides.

|  | Helix (%) | Beta (%) | Coil (%) | Turn (%) | Favored Region (%) | Allowed Region (%) | Outlier Region (%) |
| --- | --- | --- | --- | --- | --- | --- | --- |
| Epi-1 | 47 | - | 52 | 19 | 84.2 | 10.5 | 5.3 |
| Var-1 | 76 | - | 23 | - | 94.7 | 5.3 | - |
| Var-2 | 76 | - | 23 | 19 | 89.5 | 10.5 | - |

Var-1 has no beta turn, whereas epi-1 and var-2 have 19% beta turn. For epi-1, 94.7% falls in the favored region, while for var-1 and var-2, 84.2% and 89.5% fall in the favored region, respectively. Similarly, epi-1 and var-2 have 10.5% in the allowed region, and var-1 has 5.3% in the allowed region. None of the variants fall in the outlier region, while epi-1 has 5.3% in the outlier region. For the variants, the helicity increased, and the random coil content decreased. Additionally, the transition of amino acids to the favored region increased for the variants.

### 3.3. Bacteriostatic effect of epi-1 and its variants

The antimicrobial activity of the AMPs against bacterial pathogens were quantitated by measuring the growth of the pathogens with and without peptides. Absorbance (Optical Density) at 595 nm was used to measure the growth. Gram negative *E. coli* (MTCC 2939), *H. influenzae* (ATCC 25922), *K. pneumoniae* (MTCC 432), *P. aeruginosa* (MTCC 741) and Gram-positive *S. pneumoniae* (ATCC 49619) and *S. aureus* (ATCC 25923) were treated with the AMPs at concentrations 1, 2, 4, 8, 16, 32, 64, 128, 256, and 512 µg ml^-1^ to determine their MIC. Figure 4 shows the antimicrobial activity of the peptides (epi-1, var-1 and var-2) inhibiting the growth of the pathogens as bar graphs.

**Figure 4.**
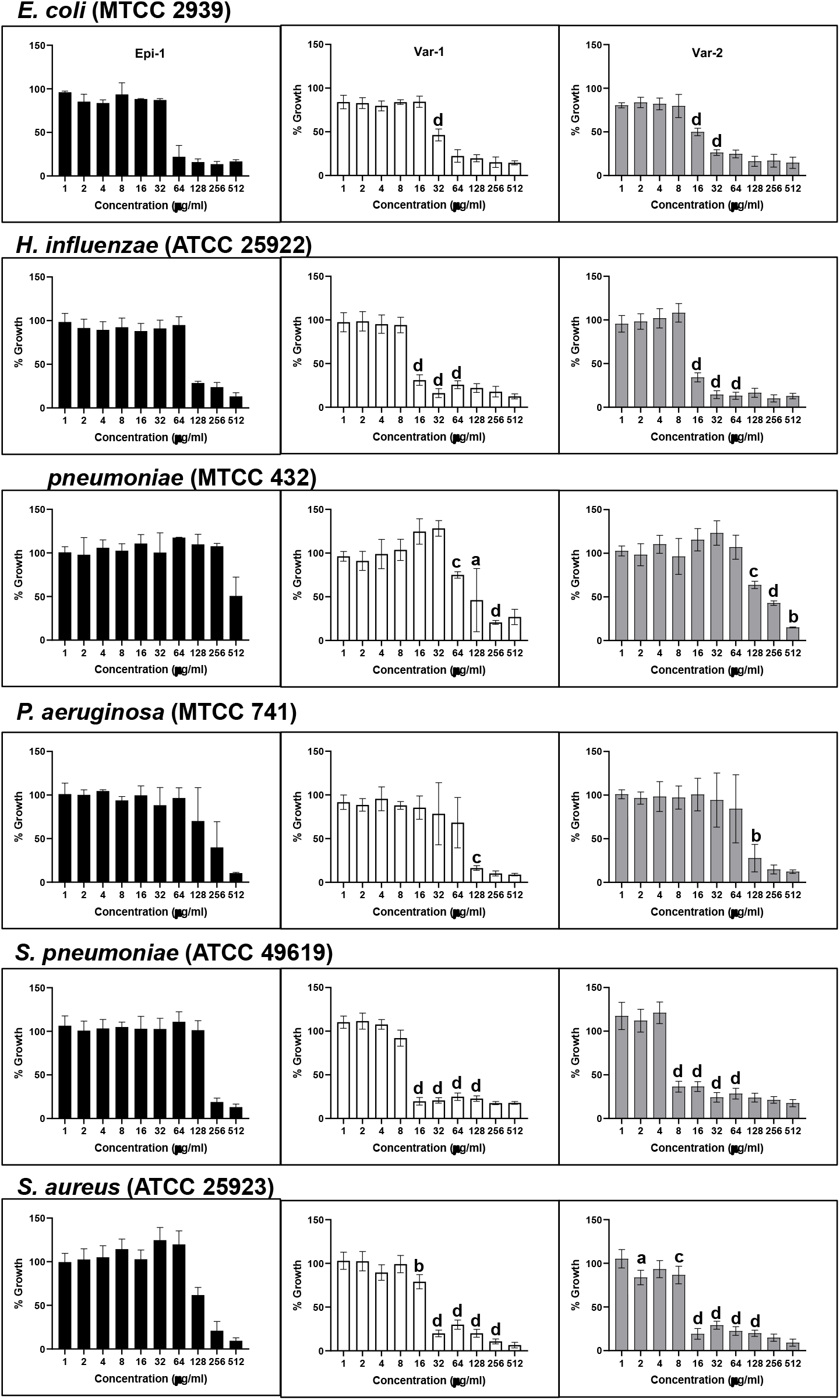
Bacteriostatic activity of epi-1 and variants against (a) *E. coli* (MTCC 2939), (b) *H. influenza* (ATCC 25922), (c) *K. pneumoniae* (MTCC 432), (d) *P. aeruginosa* (MTCC 741), (e) *P. aeruginosa* (MTCC 741), (f) *S. pneumoniae* (ATCC 49619) and (g) *S. aureus* (ATCC 25923). AMPs at concentrations 1, 2, 4, 8, 16, 32, 64, 128, 256, and 512 µg ml^-1^ were added to the pathogens in the microtitre plate and incubated for 24 h. The bars black, white and grey represent epi-1, var-1 and var-2. The growth of the organisms was measured via absorbance (OD) at 595 nm and the percentage of growth was calculated compared to control (growth without peptides). All the experiments were performed in triplicates and the standard deviation ± mean was calculated. The statistical significance representations are a; p < 0.5, b; p < 0.1, c; p < 0.01, and d; p < 0.001 comparing epi-1 vs var-1 and var-2. The photographs of the culture plates are shown in Supplementary Figure 2 and 3.

**Figure 5.**
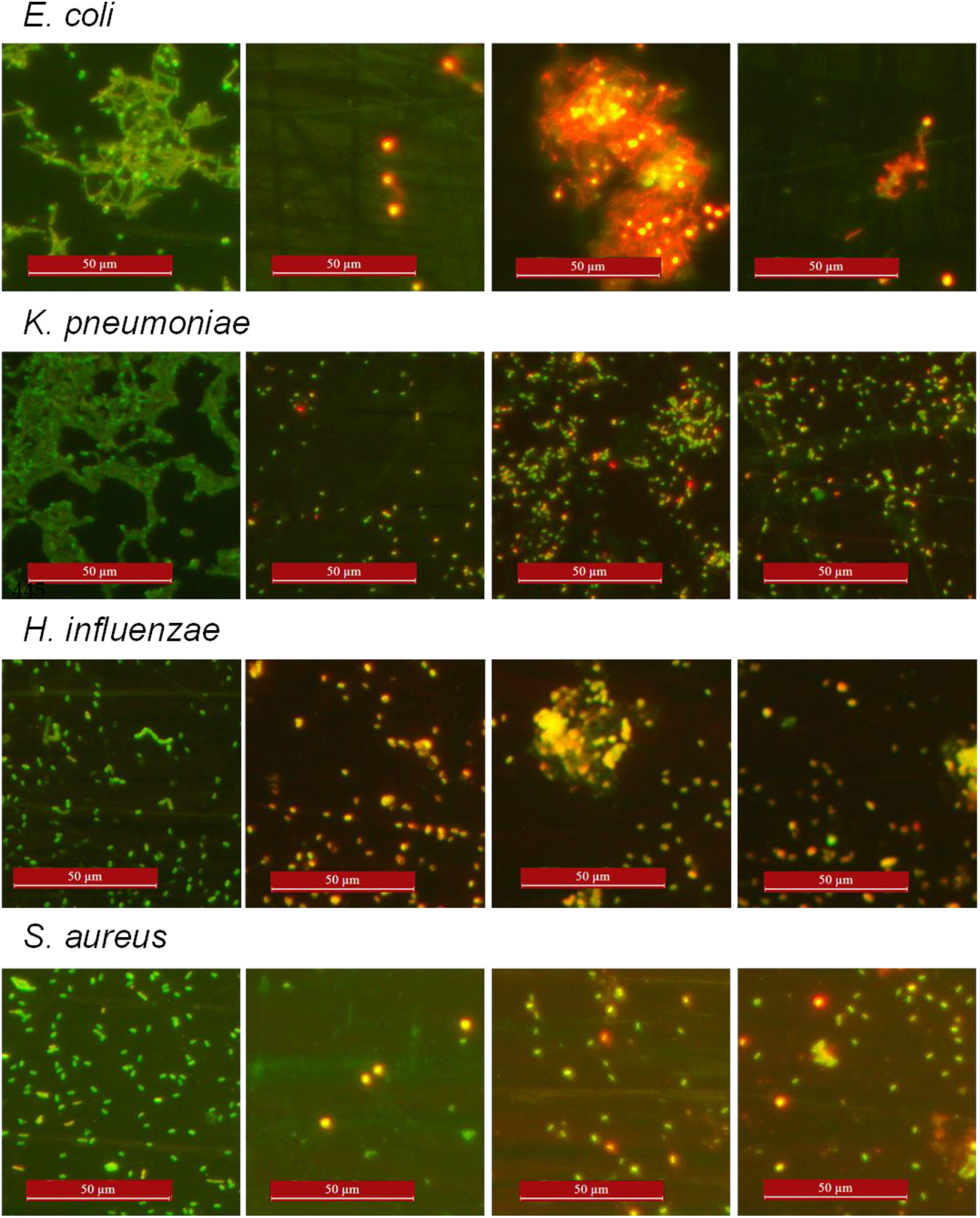
Gram-negative *E. coli, K. pneumoniae* and *P. aeruginosa* and Gram-positive *S. aureus* were stained with acridine orange and ethidium bromide and observed under a fluorescence microscope at 40X magnification. First vertical left panel: control cells. Second, third and fourth vertical panels represent epi-1, var-1 and var-2 treated at their MIC for 6h. The control pathogenic bacteria not treated with peptides have intact cell wall so they were accessible only to acridine orange but not to ethidium bromide hence appear green. Peptide-treated cells exhibited more red spots (ethidium bromide staining the genomic DNA), indicating that AMP treatment created pores in the cell walls. The peptide treated cells were entirely stained red or orange-yellow. In the case of variant peptide-treated aggregates of *E. coli* and *H. influenzae* are seen. For other bacteria, *P. aeruginosa* and *S. aureus*, fewer cells were observed in the field, stained orange or red.

Table 4 presents the minimum inhibitory concentrations (MIC) of peptides against the pathogens studied, as determined by the growth inhibition data shown in the bar graphs. In the Gram-negative category, *E. coli* has MIC values of 64 µg ml^-1^ for both epi-1 and var-1, and 32 µg ml^-1^ for var-2. There is twofold difference in activity enhancement between epi-1 vs var-2 and var-1 vs var-2. For *H. influenzae*, the MIC value is 128 µg ml^-1^ for epi-1 and 16 µg ml^-1^ for the variants. The fold difference between epi-1 and both variants (var-1 and var-2) is 8. Additionally, *K. pneumoniae* has MIC values for epi-1 and var-2 is 512 µg ml^-1^ and for var-1 it is 256 µg ml^-1^. A two-fold difference is observed between epi-1 and var-1, as well as between var-1 and var-2. *P. aeruginosa* shows MIC values of 512 µg ml^-1^ for epi-1 and 128 µg ml^-1^ for both variants. The fold difference between epi-1 and both variants (var-1 and var-2) is 4.

**Table 4.**
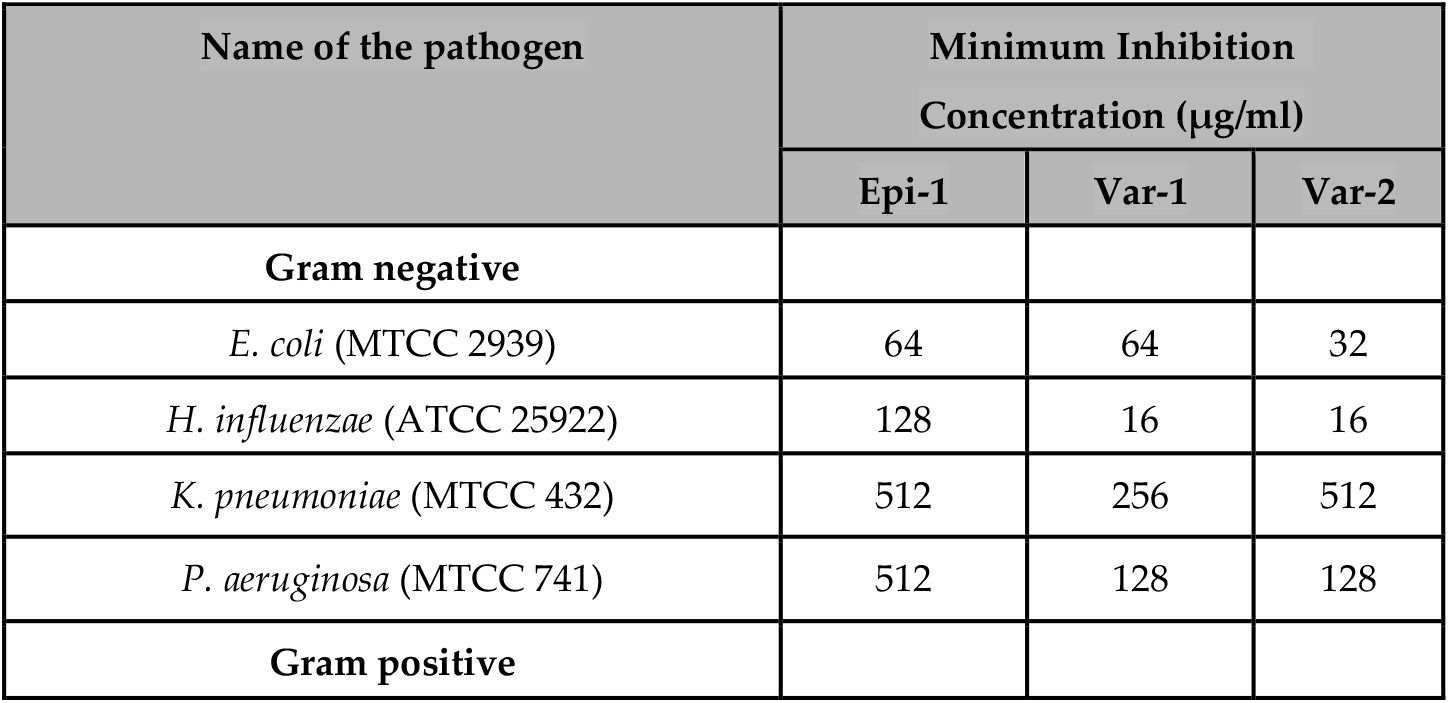
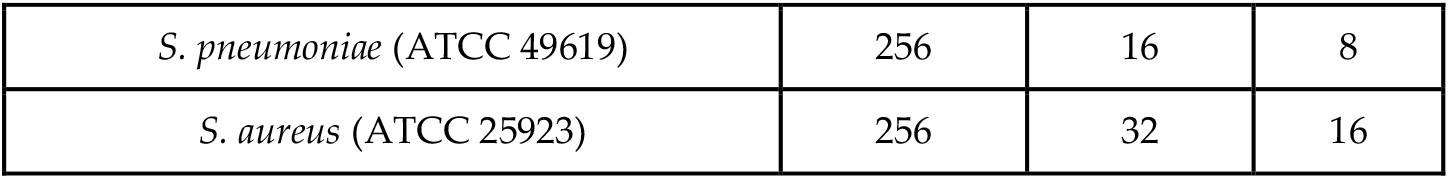
MIC values for the antimicrobial activity of epinecidin-1, variant-1 and variant-2 against pathogens. Turbidometric measurement was done after 16 hours of incubation with peptides in Müller Hington broth.

**Table 5.**
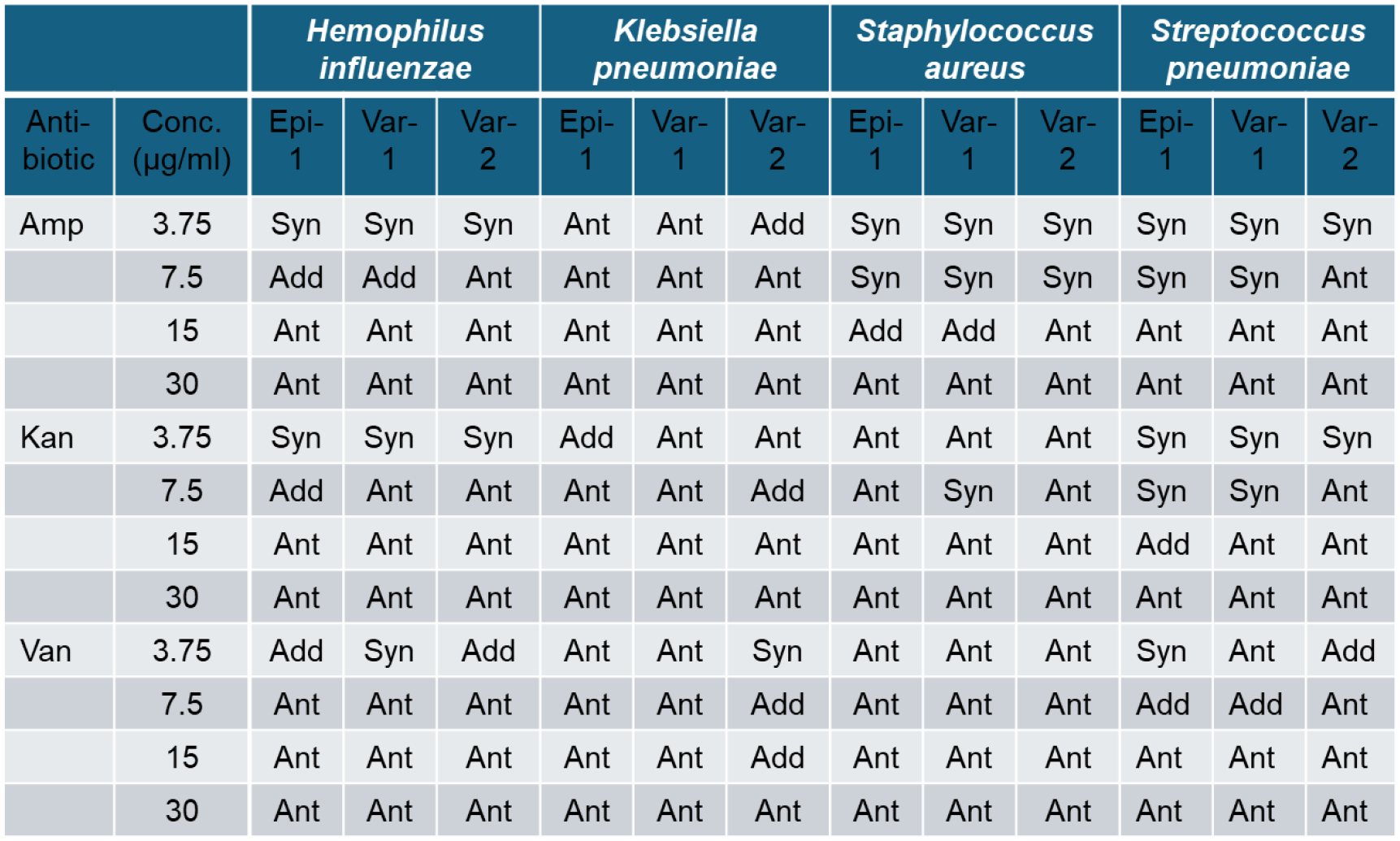
Loewe Additivity Analysis of Antibiotic and Peptide Combinations.The combination index (CI) was determined for epinecidin-1 and its variants in combination with the non-peptide antibiotics ampicillin, kanamycin, and vancomycin across a concentration range of 3.5 µg/mL to 30 µg/mL. Each data point represents an equimolar mixture of the antibiotic and the antimicrobial peptide. CI values were interpreted as follows: synergism (CI < 1), additivity (CI = 1), and antagonism (CI > 1). Synergistic and additive interactions were predominantly observed at lower concentrations. The corresponding antimicrobial activities of individual agents and their combinations are presented in Supplemental Figures 4-7.

For Gram-positive bacteria, the MIC values for *S. pneumoniae* are 256 µg ml^-1^ for epi-1, 16 µg ml^-1^ for var-1, and 8 µg ml^-1^ for var-2. The fold difference between epi-1 and var-1 is 16, between epi-1 and var-2 is 32, and between var-1 and var-2 is 2. Similarly, for *S. aureus*, the MIC values are 256 µg ml^-1^ for epi-1, 32 µg ml^-1^ for var-1, and 16 µg ml^-1^ for var-2.

These results denote that all the peptides have bacteriostatic activity against the tested Gram-negative and Gram-positive bacterial pathogens, however, the degree of activity varies and overall, the variants show better antimicrobial activity than epi-1. The fold difference between epi-1 and var-1 is 8, between epi-1 and var-2 is 16, and between var-1 and var-2 it is 2.

### 3.4. Impact of epi-1 and variants on pathogenic bacteria as observed through AO/EtBr differential staining

AO/EtBr differential staining is a sensitive technique for screening live and dead cells after peptide treatment. The superiority of EtBr fluorescence over AO indicates significant cell death. *E. coli* cells treated with peptides absorb EtBr and appear red, demonstrating rapid cell death. For other bacterial strains, yellow-orange (due to mix of green and red) to red fluorescence is observed demonstrating significant membrane damage and cell death. Control cells appear green due to acridine orange staining, as they resist EtBr entry.

### 3.5. Combination effect of epi-1 variants with non peptide antibiotics on antimicrobial activity

The combination effects of three antibiotics ampicillin, kanamycin and vancomycin at varying concentrations (3.75, 7.5, 15, and 30 µg/ml) with AMPs at the same concentrations were tested against four bacterial species: *Hemophilus influenzae, Klebsiella pneumoniae, Staphylococcus aureus*, and *Streptococcus pneumoniae*.

For *Hemophilus influenzae*, ampicillin and kanamycin show synergistic effects at low concentrations but become antagonistic at higher concentrations. For *Klebsiella pneumoniae*, kanamycin consistently shows additive effects across all strains and concentrations, while ampicillin is mostly antagonistic. For *Staphylococcus aureus*, ampicillin shows synergistic or additive effects at lower concentrations but becomes antagonistic at higher levels. For *Streptococcus pneumoniae*, all antibiotics consistently show synergistic effects across all concentrations and strains.

### 3.6. Epinecidin-1 and its variants accelerated wound healing activity in rat epidermis

Wound healing efficacy of the peptides was assessed using a full-thickness excisional wound model in rats. Circular wounds (5 mm in diameter) were created using a biopsy punch, and morphological changes were monitored over an 8-day period. Peptides were topically applied once daily, while phosphate-buffered saline (PBS) served as the vehicle control. Figure 6a presents the macroscopic appearance of wounds following treatment with epinecidin-1 and its variants. Visual inspection revealed accelerated wound closure in peptide-treated groups, characterized by enhanced connective tissue formation and reduced wound area compared to controls. By day 8, peptide-treated wounds exhibited near-complete filling with granulation tissue, whereas control wounds remained partially open. Following euthanasia on day 8, wound tissues were harvested, sectioned, and stained with Hematoxylin and Eosin (H&E) for histological evaluation.

**Figure 6.**
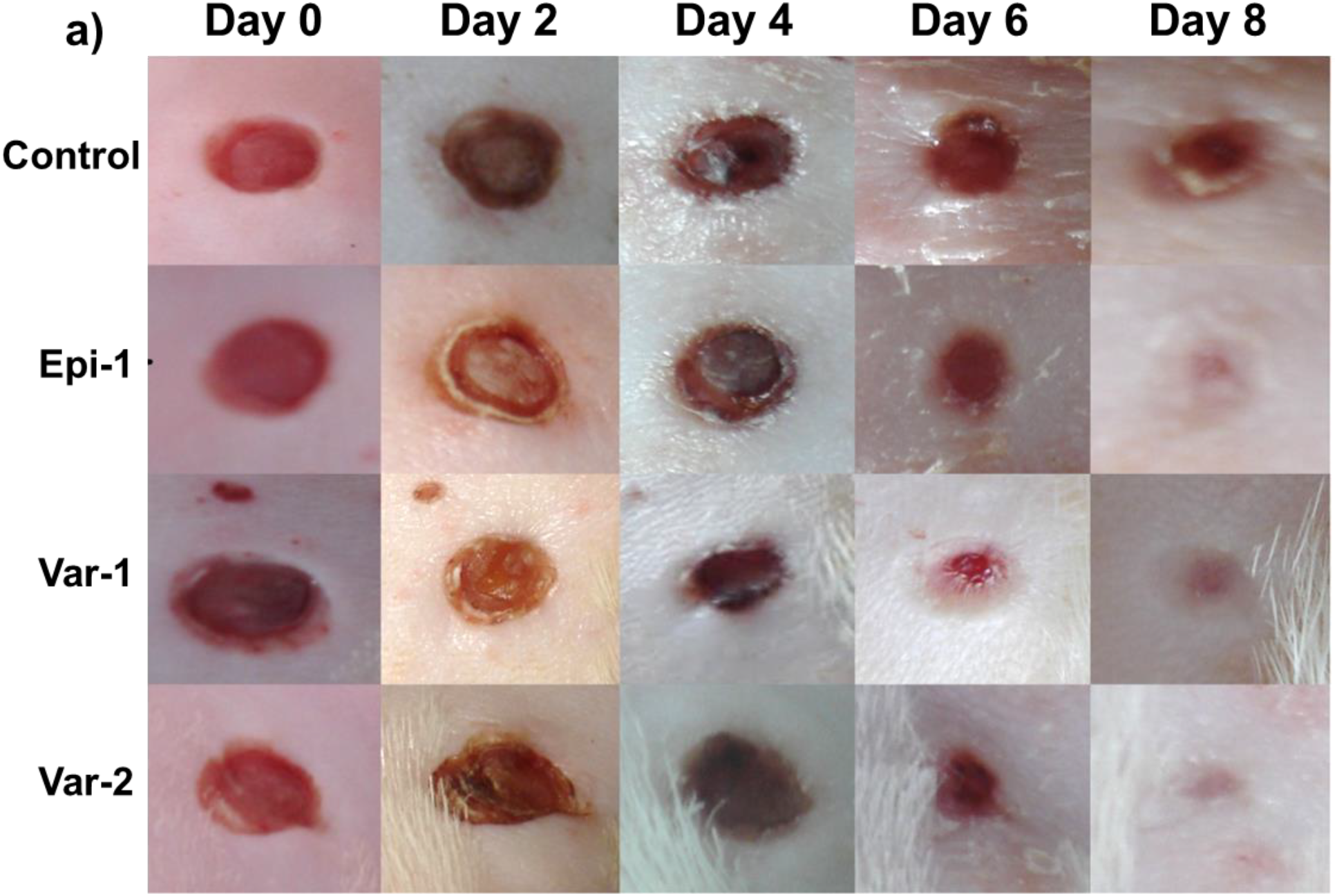
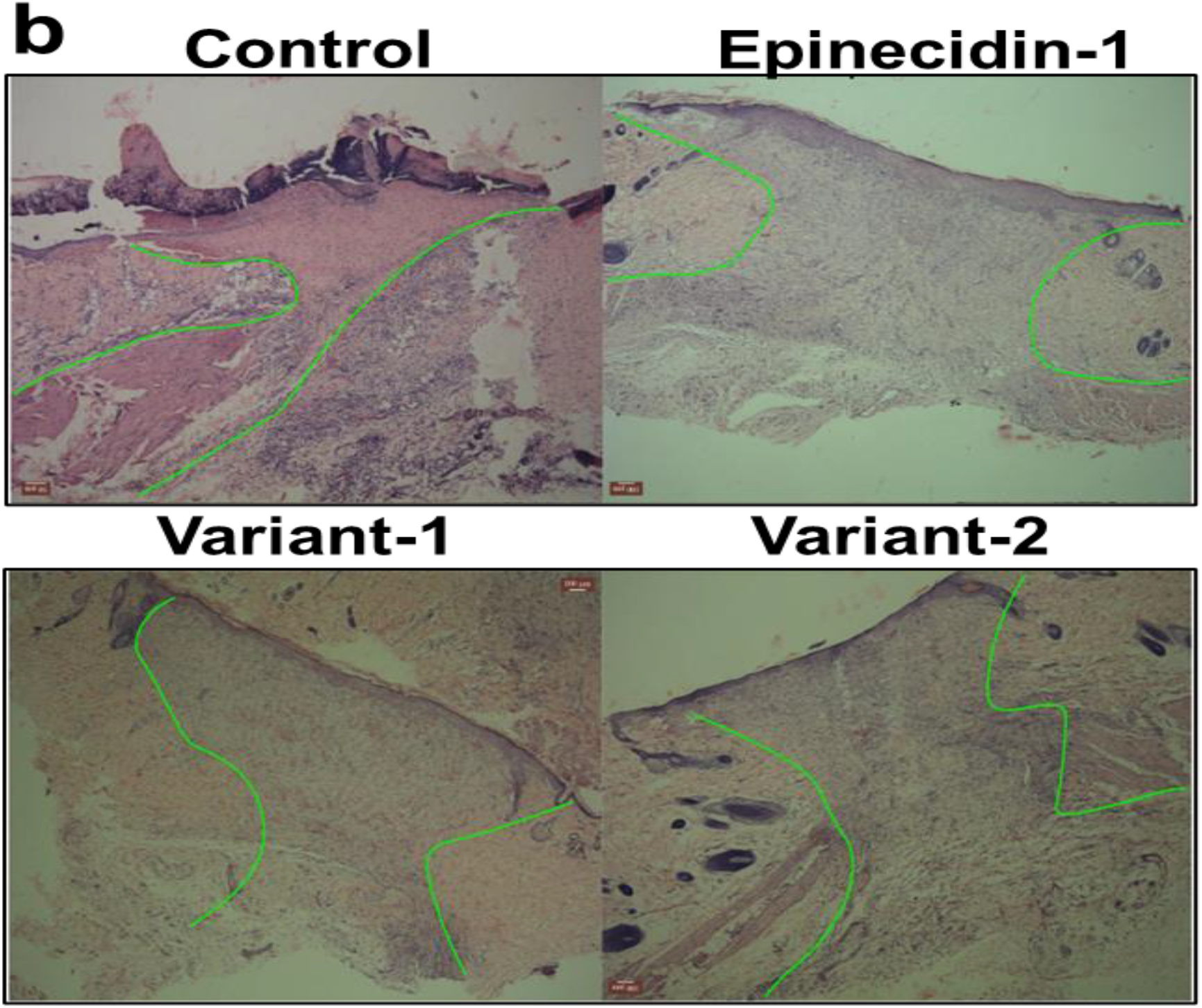
Epinecidin-1 enhanced wound healing in a murine model. (a) Representative macroscopic images showing wound closure progression over time in peptide-treated versus vehicle control groups. Peptide treatment resulted in visibly accelerated wound contraction. (b) Histological analysis of wound cross-sections harvested on day 8 post-injury and stained with Hematoxylin and Eosin (H&E). Peptide-treated tissues exhibited increased granulation tissue formation (highlighted between green borders), indicating enhanced tissue regeneration compared to controls.

Figure 6b shows representative micrographs of wound cross-sections. Peptide-treated tissues displayed significantly increased granulation tissue, with a higher density of granulocytes, indicating robust inflammatory and proliferative responses. Furthermore, re-epithelialization was markedly improved in peptide-treated wounds, as evidenced by the nearly continuous keratinocyte layer, in contrast to the incomplete epithelial coverage observed in controls. Notably, a substantial increase in neovascularization was observed in the peptide-treated groups, with numerous newly formed blood vessels visible within the granulation tissue (highlighted between green borders), suggesting that epinecidin-1 and its variants strongly promote angiogenesis *in vivo*. Collectively, these findings underscore the therapeutic potential of these peptides in enhancing wound healing.

## 4. Discussion

Administering new antibiotics or a combination therapy with multiple drugs is necessary when dealing with infections caused by antibiotic-resistant bacteria [31]. Such combined therapy not only reduces the high drug dosage but also decreases the chances of development of resistance in bacteria. These short peptides aggregates the bacterial proteins or other proteins it comes in contact with [32-35]. The short cationic peptides also bind to DNA [36, 37]. High-throughput research has developed synthetic AMPs with modified amino acid sequence, effective against a vast range of pathogens. These aspects have rekindled interest in exploring the synergistic effects in antibiotic actions in numerous research laboratories.

The antimicrobial peptide Epinecidin-1 has demonstrated significant potential in disrupting biofilms [18-20, 38-43], which are typically resistant to conventional antibiotics due to their protective extracellular matrix and reduced permeability. This activity is largely attributed to Epinecidin-1’s membrane-penetrating properties. In this study, we investigated its synergistic effects when combined with conventional antibiotics. Structural prediction of epinecidin-1 and its variants using Ramachandran plot reveals that the peptide variants exhibit enhanced helicity and conformational stability compared to the native epinecidin-1. Notably, the variants (Var-1 and Var-2) display a helical content of 76%, in contrast to 47% observed in Epi-1. Structural and bio-physical characterization using RP-HPLC co-relates with the predicted characteristics. The Variant-1 is a perfect amphipathic peptide. Hydrophobic on one surface and hydrophilic (cationic) on the opposite surface.

Previously, we investigated the *in vitro* activity of recombinant purified epi-1 and its variants against nosocomial pathogens obtained from repositories and clinical locations[18-20, 41]. In this study we explored the potential of combining these peptides with conventional antibiotics such as ampicillin and vancomycin by Leowe’s combination principle. For *Haemophilus influenzae*, ampicillin and kanamycin exhibit synergistic interactions at lower range of concentrations tested with all the three AMPs. In this synergistic effect, two fold less amount of peptide and antibiotics were used to achieve the same effect when used alone. In combination with the peptides, even more effective pathogen killing was observed. For *Klebsiella pneumoniae*, kanamycin consistently demonstrates additive effects across all tested strains and concentrations, whereas ampicillin predominantly shows antagonistic behavior. For *Staphylococcus aureus*, ampicillin shows synergistic or additive effects at low concentrations but becomes antagonistic as the concentration increases. For *Streptococcus pneumoniae*, all the three tested antibiotics show consistent synergistic effects against concentration range. The variability observed in combination efficacy among strains studied may plausibly be attributed to differences in bacterial cell surface properties, which serve as primary interaction sites for antimicrobial peptides (AMPs) [44]. Additionally, the distinct mechanisms of action associated with each antibiotic likely contribute to the differential synergistic outcome [45].

The limitation of this study is that we did not evaluate a wide range of concentration. However the preliminary tests revealed that epinecidin-1 and its variants exhibited synergistic and additive interactions with the three antibiotics at specific concentrations. The reasons behind these variable effects observed between the AMPs and antibiotics remain unclear. In some concentrations used, the combination with antibiotics enhances antimicrobial activity, while some concentrations does not. To better understand this phenomenon, further studies involving a broader range of bacterial strains and detailed molecular-level investigations are needed. Although it has been suggested that the antibiotic resistance phenotype might influence these outcomes, existing research indicates that AMP efficacy is generally independent of bacterial resistance profiles [46]. This combination approach could enhance the effectiveness of traditional antibiotics, making it a promising strategy for managing bacterial infections. These results provide new data that can be used in developing new treatments for multidrug-resistant bacterial infections.

In addition to their antimicrobial activity, the peptides were assessed for their wound healing efficacy *in vivo* using a rat model. The results indicated that epinecidin-1 and its variants significantly accelerated the healing process. In Figure 6, there is evidence for epithelial cell migration in peptide treated wounds. The AMPs interactions *in-vivo* would have been due to selective homing to angiogenic vessels of wound tissue. Thus the interaction with the wound may cause fibronectin deposition leading to migration of epidermis to close the wound. This addresses the treatment of skin wounds also.

The findings suggest that epinecidin-1 and its variants can be used with antibiotics for synergistic activity in clinical and industrial applications. The short AMPs are not only effective antimicrobial agents but also have the potential to improve wound healing and enhance the efficacy of existing antibiotics. This dual functionality makes them strong candidates for clinical and industrial applications, particularly in managing bacterial infections in both human and livestock populations.

## 5. Conclusion

This study provides strong evidence supporting the synergistic antimicrobial and wound-healing efficacy of epinecidin-1 and its variants with antibiotics, underscoring their potential as innovative therapeutic agents for managing nosocomial infections and enhancing clinical outcomes.

## Supplementary Materials

Supplementary Figure-1: Elutuon profile of the peptides in RP-HPLC column.

Supplementary Figure-2: Photographs of pathogens tested for MIC against *E.coli, H. influenzae, K. pneumoniae*.

Supplementary Figure-3: Photographs of pathogens tested for MIC against *P. aeruginosa, S. pneumoniae, S. aureus*.

Supplementary Figure-4: Antimicrobial activity normalized to growth of *Haemophilus influenzae* incubated each peptide and their combination with antibiotics.corresponding antimicrobial activities of individual agents and their combination

Supplementary Figure-5: Antimicrobial activity normalized to growth of *Klebsiella pneumoniae* incubated each peptide and their combination with antibiotics.corresponding antimicrobial activities of individual agents and their combination

Supplementary Figure-6: Antimicrobial activity normalized to growth of *Streptococcus pneumoniae* incubated each peptide and their combination with antibiotics.corresponding antimicrobial activities of individual agents and their combination

Supplementary Figure-7: Antimicrobial activity normalized to growth of *Staphylococcus aureus* incubated each peptide and their combination with antibiotics.corresponding antimicrobial activities of individual agents and their combination

## Author Contributions

Conceptualization: S.J., A.K.; Methodology: S.J. and A.K.; Software: S.J. and A.K.; Validation, S.J. and A.K.; Formal analysis, S.J. and A.K.; Investigation, A.K.; Resources, S.J., A.K.; Data curation, S.J. and A.K; Writing—original draft preparation, S.J. and A.K.; Writing—review and editing, S.J. R.V, A.Aand A.K.; Visualization, S.J., R. V., A.A., and A.K.; supervision, A.K.; project administration, A.K.; Funding acquisition, S.J.,and A.K. All authors have read and agreed to the published version of the manuscript.

## Acknowledgements and Funding

This research was funded by the Indian Council of Medical Research, India, for partial financial support via a sanctioned research project to the corresponding author (Ref. No. IIPSG-2024-01-01898).

**Institutional Review Board Statement**

This study adhered to Institutional Animal Ethical Committee guidelines (Ref. No. BDU/IAEC/2012/42) Bharathidasan University, Tiruchirapalli, India.

## Supplementary Files

**Supplementary Figure 1.**
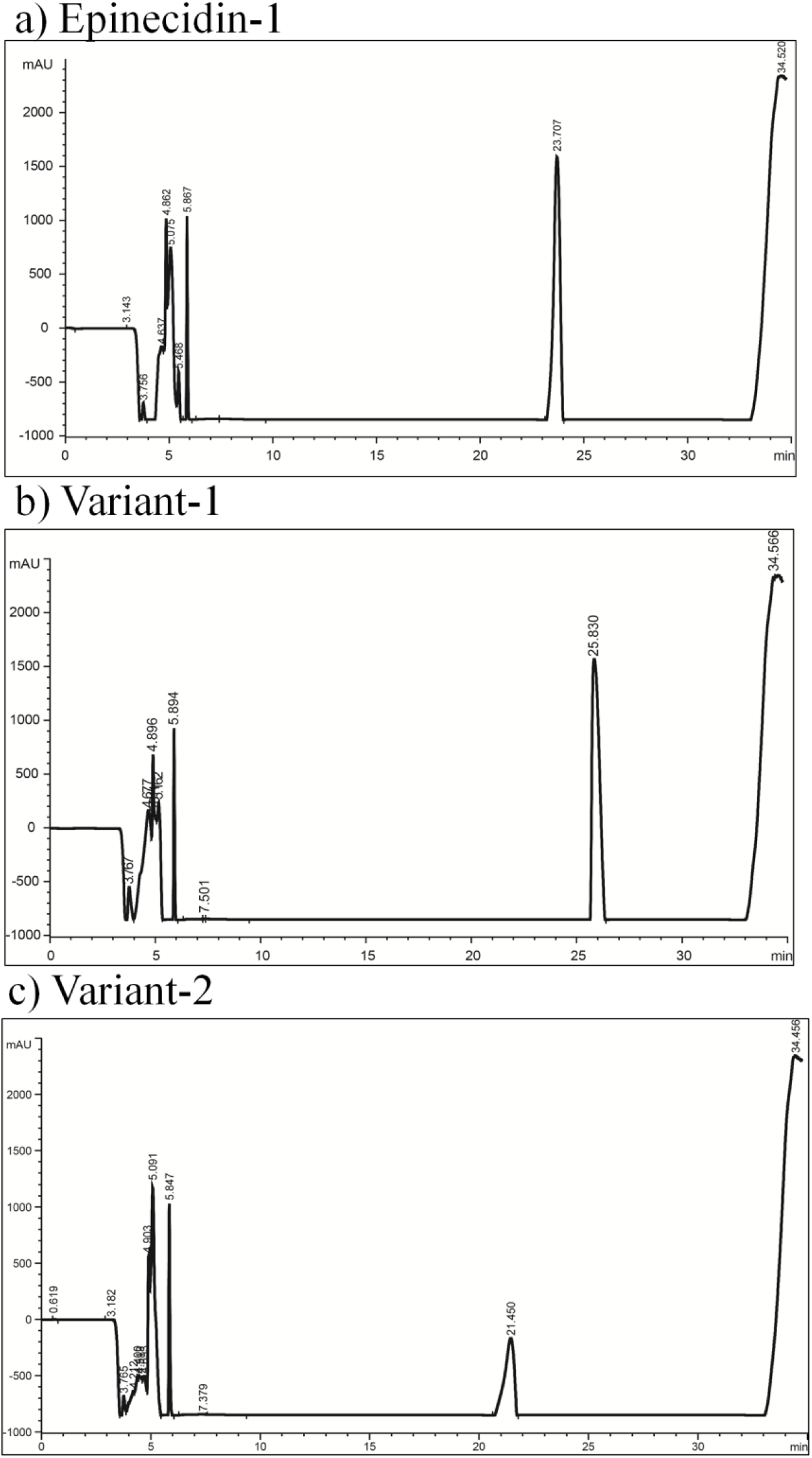
Elutuon profile of the peptides in RP-HPLC column.

**Supplementary Figure 2.**
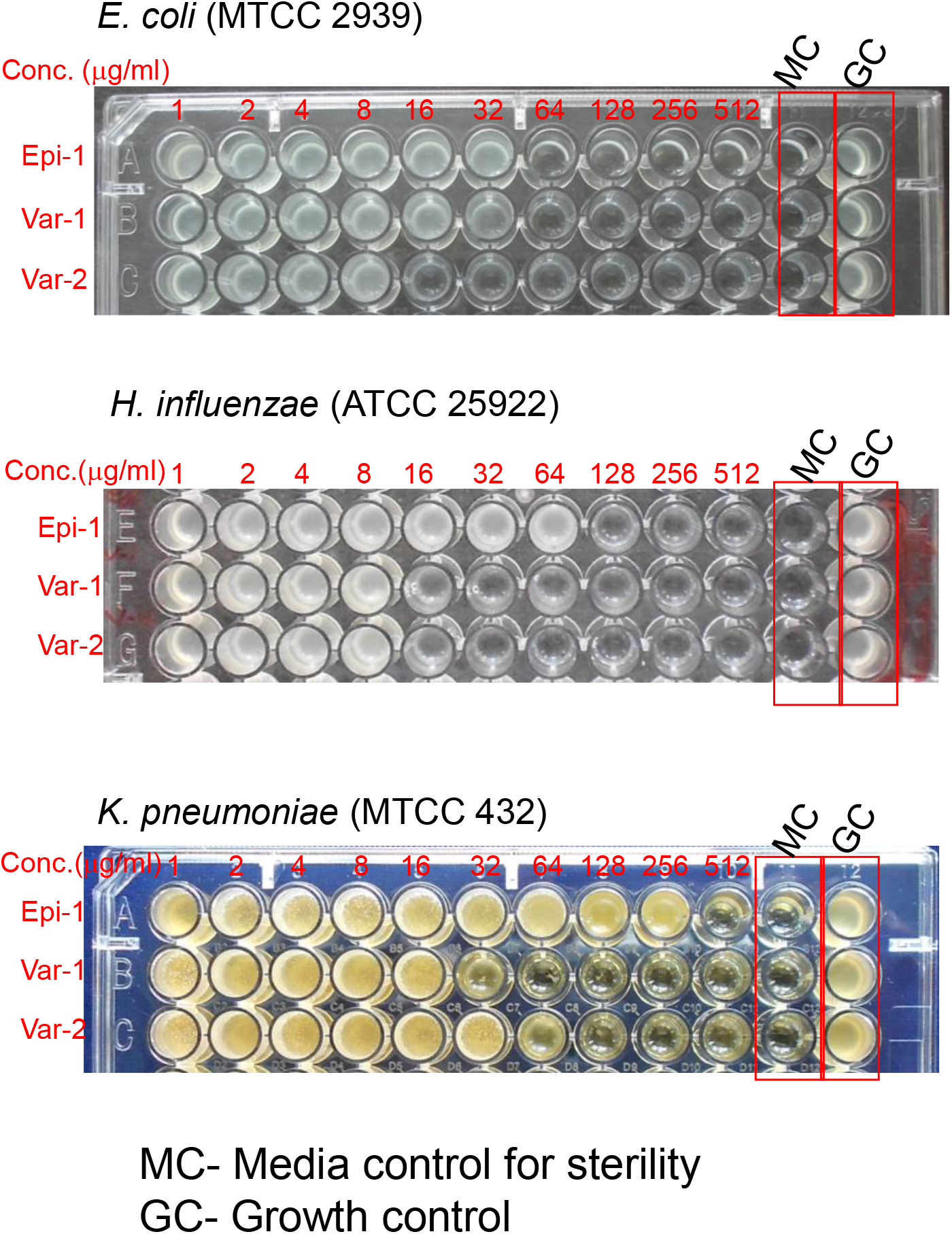
Photographs of pathogens tested for MIC.

**Supplementary Figure 3.**
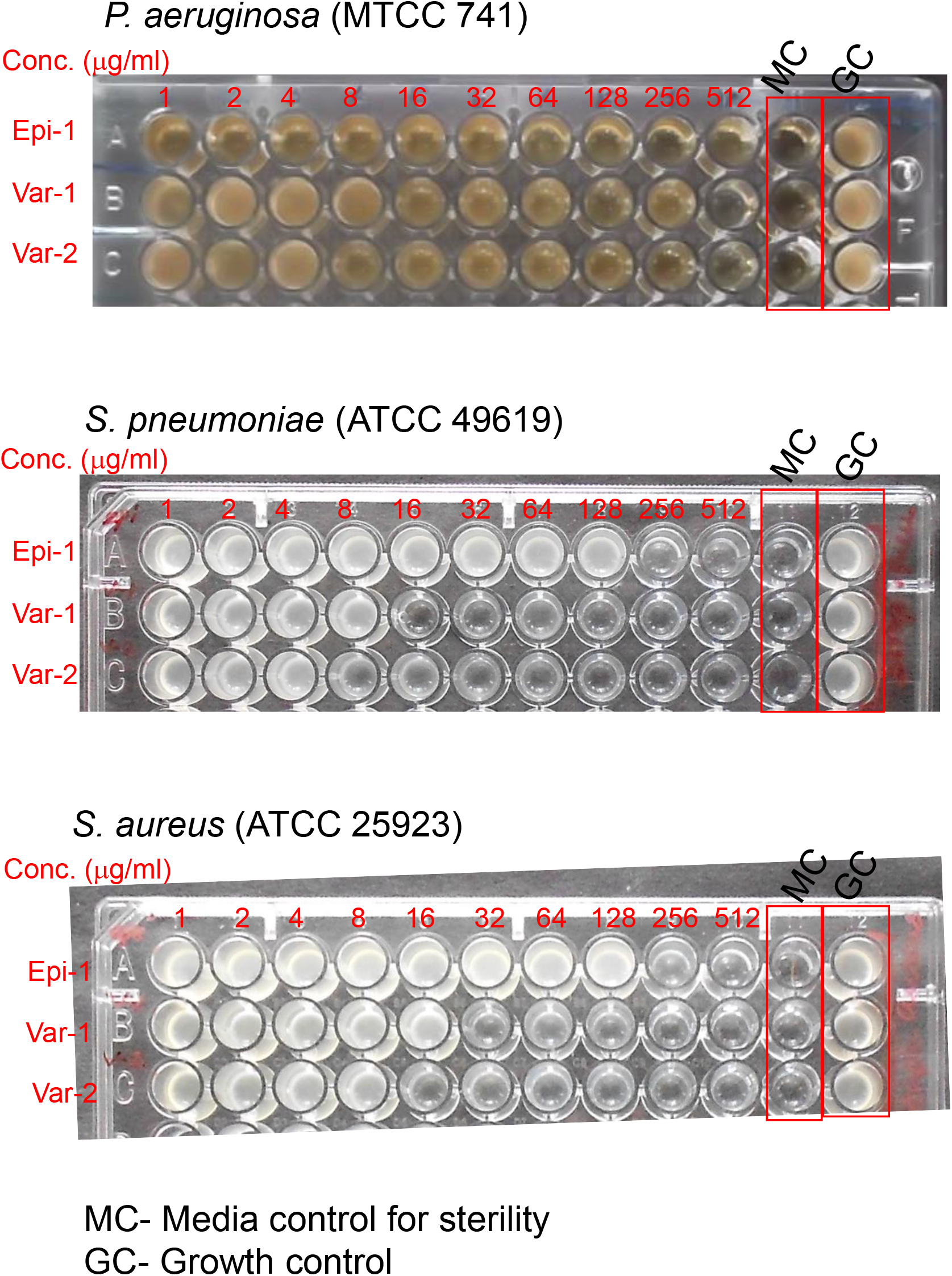
Photographs of pathogens tested for MIC.

**Supplementary Figure 4.**
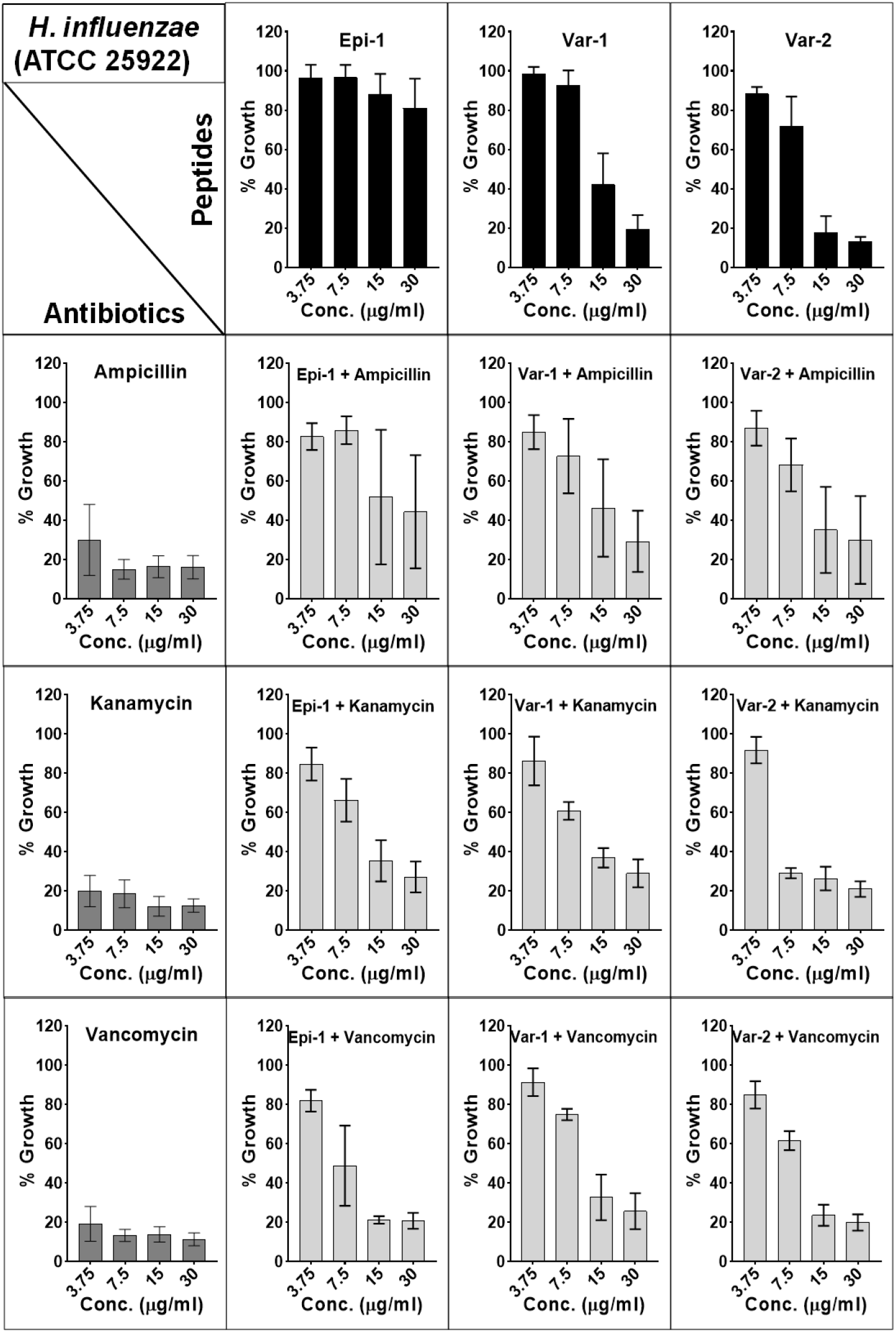
Antimicrobial activity normalized to growth of *Haemophilus influenzae* incubated each peptide and their combination with antibiotics.corresponding antimicrobial activities of individual agents and their combination.

**Supplementary Figure 5.**
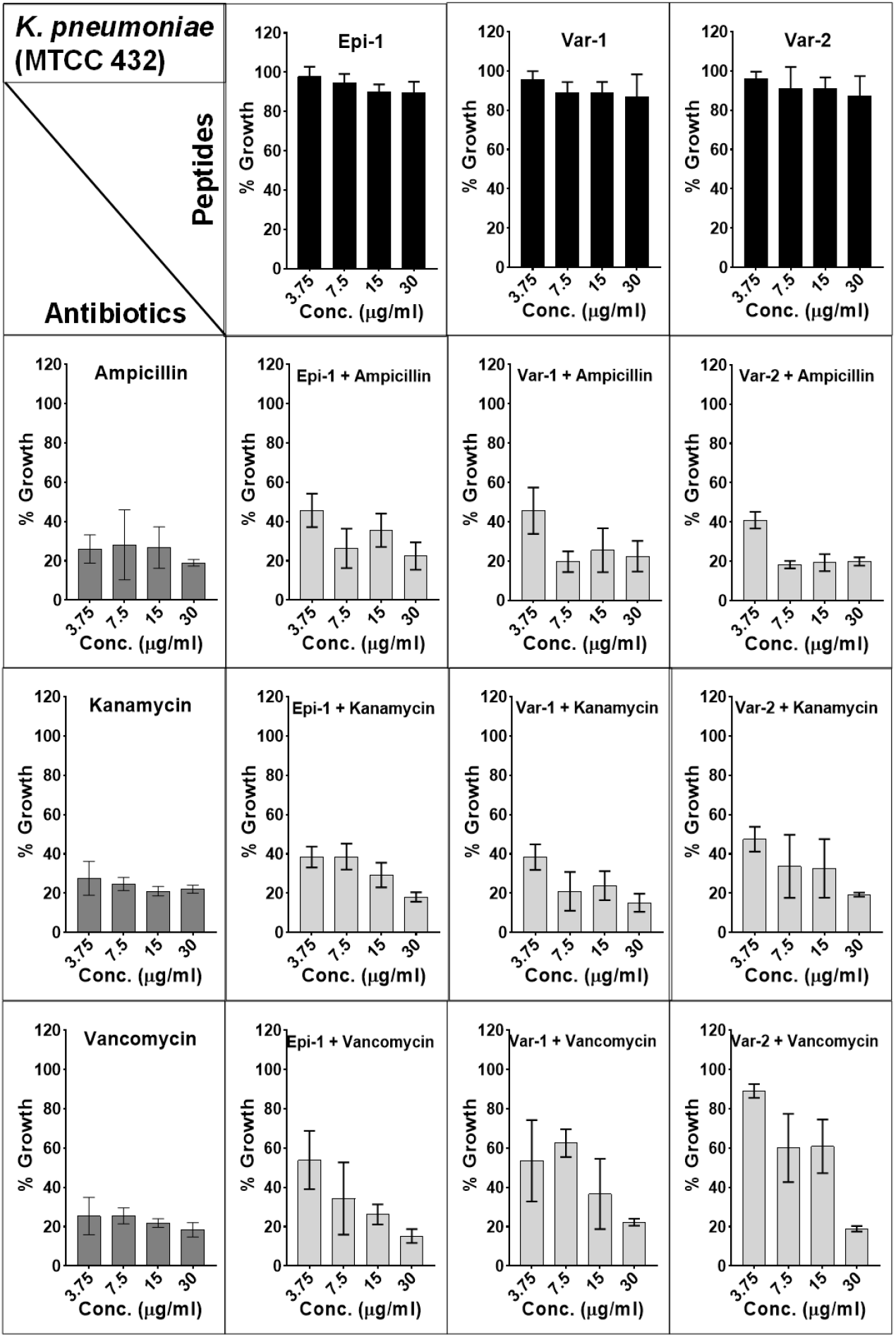
Antimicrobial activity normalized to growth of *Klebsiella pneumoniae* incubated each peptide and their combination with antibiotics.corresponding antimicrobial activities of individual agents and their combination.

**Supplementary Figure 6.**
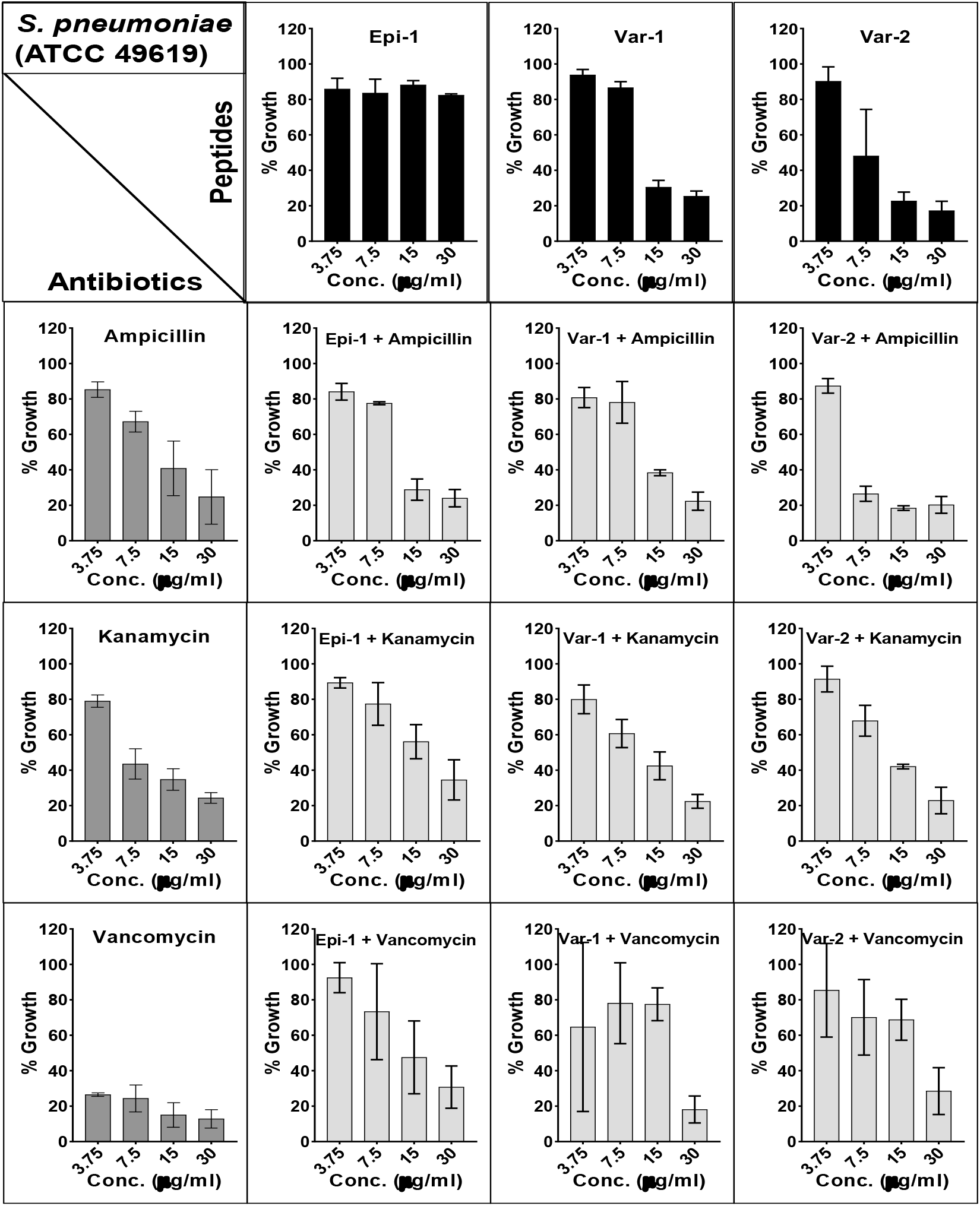
Antimicrobial activity normalized to growth of *Streptococcus pneumoniae* incubated each peptide and their combination with antibiotics.corresponding antimicrobial activities of individual agents and their combination.

**Supplementary Figure 7.**
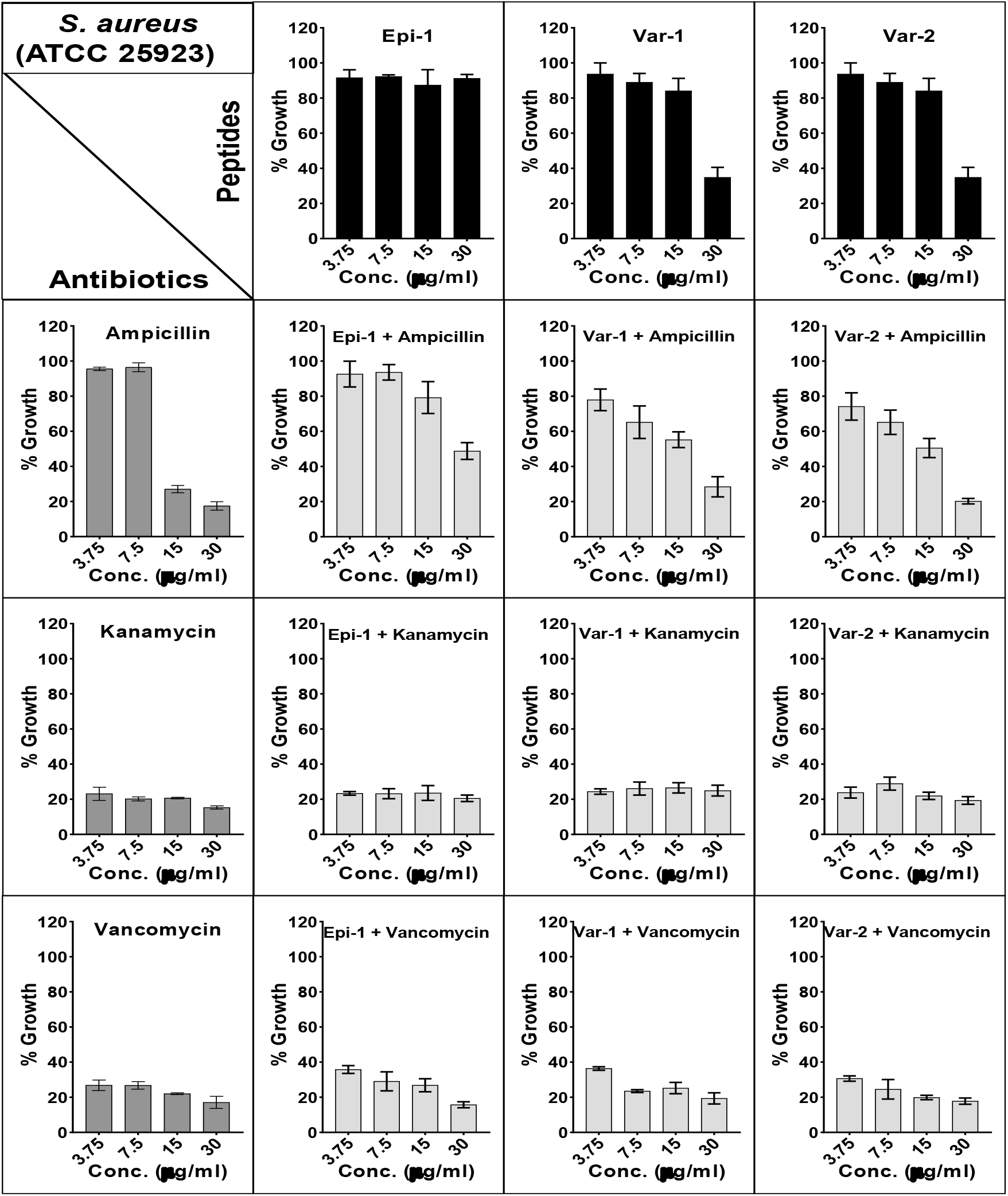
Antimicrobial activity normalized to growth of *Staphylococcus aureus* incubated each peptide and their combination with antibiotics.corresponding antimicrobial activities of individual agents and their combination.

